# The impact of stomatal kinetics on diurnal photosynthesis and water use efficiency under fluctuating light

**DOI:** 10.1101/2020.09.03.281873

**Authors:** David Eyland, Jelle van Wesemael, Tracy Lawson, Sebastien Carpentier

**Affiliations:** Laboratory of Tropical Crop Improvement, Division of Crop Biotechnics, KU Leuven, Leuven, Belgium; School of Life Sciences, University of Essex, Colchester, Essex, UK; Bioversity International, Banana Genetic Resources, Leuven, Belgium

## Abstract

Dynamic light conditions require continuous adjustments of stomatal aperture. As stomatal conductance (g_s_) kinetics are a magnitude slower than photosynthesis (*A*), they are hypothesized to be key to plant productivity and water use efficiency. Using step-changes in light intensity, we studied the diversity of light-induced g_s_ kinetics in relation to stomatal anatomy in five banana genotypes (*Musa* spp.) and modelled the impact on *A* and intrinsic water use efficiency (_i_WUE). Banana generally exhibited a strong limitation of *A* by g_s_, indicating a priority for water saving. Significant genotypic differences in g_s_ kinetics and g_s_-based limitations of *A* were observed. For two contrasting genotypes the impact of differential g_s_ kinetics on *A* and _i_WUE was further investigated under realistic diurnally fluctuating light conditions and at whole-plant level. Genotype-specific stomatal kinetics observed at the leaf level were corroborated at whole-plant level, suggesting that despite differences in g_s_ control at different locations in the leaf and across leaves, genotype-specific responses are still maintained. However, under diurnally fluctuating light conditions g_s_ speediness had only a momentary impact on the diurnal _i_WUE and carbon gain. During the afternoon there was a setback in kinetics: the absolute g_s_ and the g_s_ responses to light were damped, strongly limiting *A* and the diurnal _i_WUE. We conclude that the impact of the differential g_s_ kinetics on the limitation of *A* was dependent on the target light intensity, the magnitude of change, the g_s_ prior to the intensity change and particularly the time of the day.

**One sentence summary:** Genotype-specific stomatal rapidity is for the first time validated at whole-plant level, but under fluctuating light the impact of stomatal dynamics depends on other factors like the time of the day.

## Introduction

In order to survive, plants need to balance CO_2_ uptake for photosynthesis with water loss via transpiration. By adjusting their aperture, stomata control gaseous exchange between the leaf interior and the external atmosphere. Stomatal aperture is adjusted by moving solutes into or out of the guard cells. These changes in osmotic potential elicit water movement in or out of the guard cells, altering turgor pressure and subsequently aperture. In general, stomatal opening in well-watered C3 and C4 species is triggered by high light intensity, low VPD and low CO_2_ concentrations. Opposite environmental conditions (low light, high VPD and high [CO_2_]) stimulate stomatal closure (Assmann and Shimazaki, 1999; Outlaw, 2003; Lawson and Morison, 2004). Therefore, in a dynamic field environment, stomata are continuously adjusting aperture to achieve an appropriate balance between carbon gain and water loss (Pearcy, 1990; Lawson and Blatt, 2014). However, most research has studied stomatal conductance (g_s_) and photosynthesis (*A*) under steady-state conditions. A high g_s_ under steady-state conditions is associated with high *A* and consequently improved growth (Fischer et al., 1998; Franks, 2006). However, as g_s_ kinetics are a magnitude slower than those of *A*, the speed in which these steady-state values are reached in a fluctuating environment have a great influence on the growth and water use efficiency (Lawson and Blatt, 2014; Kaiser et al., 2016; McAusland et al., 2016; Taylor and Long, 2017; De Souza et al., 2020; Yamori et al., 2020). In a fluctuating field environment, light intensity is one of the most variable environmental conditions as it changes continuously by moving cloud covers and shading from adjacent plants (Pearcy, 1990; Slattery et al., 2018; Morales and Kaiser, 2020). In this way, stomata frequently experience alternating light intensities, inducing stomatal responses that change *A*, g_s_ and the ratio of these, the intrinsic water use efficiency (_i_WUE). The balance between CO_2_ gain and H_2_O loss under changing light intensities is disturbed by delayed g_s_ responses (Vialet-Chabrand et al., 2017; Slattery et al., 2018). The slower g_s_ increase to increased light intensity limits the CO_2_ uptake for *A*, while the slower g_s_ decrease to decreased light intensity results in unnecessary water loss. The limitation of *A* by the slower kinetics of g_s_ has been shown to be significant in well-watered C3 species (Farquhar and Sharkey, 1982; Jones, 1998; Lawson and Blatt, 2014; McAusland et al., 2016). Rapid g_s_ kinetics therefore have been hypothesized to maximize *A* and _i_WUE, as steady-state values under the new conditions can be rapidly achieved (Lawson and Blatt, 2014; Papanatsiou et al., 2019; Kimura et al., 2020; De Souza et al., 2020). The g_s_ kinetics are, together with the final steady-state g_s_ the plant reaches, crucial to determine the plant performance (Franks and Farquhar, 2007; Vico et al., 2011; McAusland et al., 2016; Qu et al., 2016; Faralli et al., 2019b; Yamori et al., 2020). The importance of diversity in g_s_ kinetics under fluctuating light was highlighted by De Souza et al. (2020), who showed a three fold higher variability in carbon assimilation between cassava genotypes than under steady-state conditions, mainly caused by differences in stomatal limitation. However, to our knowledge, the diversity of g_s_ kinetics across varieties has never been investigated at whole-plant level nor under diurnally fluctuating light conditions.

Here our research aimed to explore biodiversity in light-induced stomatal dynamics across genotypes and evaluate for the first time the impact on whole-plant level. We studied the diversity of light-induced g_s_ kinetics in relation to stomatal anatomy in five banana genotypes *(Musa* spp.) and modelled the impact on *A* and intrinsic water use efficiency (_i_WUE) under both single step-changes in light intensity and realistic diurnal fluctuating light conditions. By comparing the g_s_ kinetics in response to step-changes with the g_s_ responses under fluctuating light conditions, we gain insight in the importance of stomatal kinetics on diurnal carbon gain and water use efficiency.

## Results

### *A* and g_s_ response to step-changes

Increasing light intensity from 100 to 1000 μmol m^-2^ s^-1^ induced a strong stomatal opening response (Fig. 1). The g_s_ response followed a sigmoidal pattern. A similar sigmoidal limiting pattern was observed for *A* in all genotypes, indicating a strong limitation of *A* by g_s_ in *Musa* (Fig. 1). Between genotypes there were significant differences in the speed of g_s_ increase. Steady-state *A* and g_s_ under high light intensity were reached in three out of five genotypes. In contrast, the genotype Cachaco and Leite continued to increase g_s_ and *A* slowly after 90 min of 1000 μmol m^-2^ s^-1^. The subsequent decrease in light intensity from 1000 to 100 μmol m^-2^ s^-1^ resulted in a rapid g_s_ decrease, which also followed a sigmoidal pattern (Fig. 1). Photosynthesis, on the other hand, as expected decreased instantly because light became the limiting factor (Fig. 1).

**Fig. 1.**
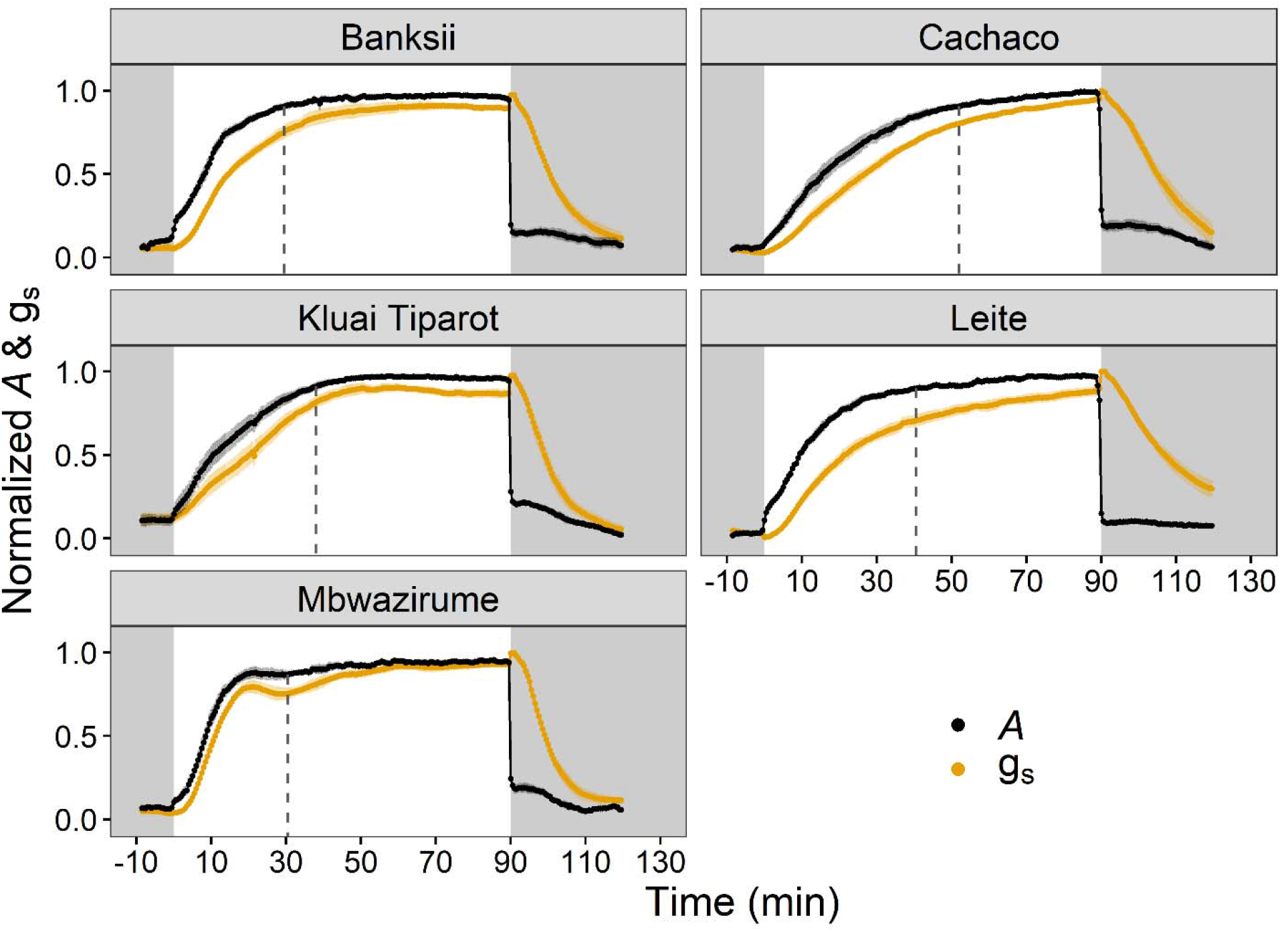
Normalized response of stomatal conductance (g_s_, orange) and photosynthesis (*A*, black) of five banana genotypes to a step increase in light intensity from 100 to 1000 μmol m^-2^ s^-1^ followed by a decrease from 1000 to 100 μmol m^-2^ s^-1^. Grey and white areas indicate time periods of 100 μmol m^-2^ s^-1^ and 1000 μmol m^-2^ s^-1^, respectively. Dashed lines indicate when 95 % of steady-state *A* was reached. Points and error bars represent mean ± SE (n = 7-8).

### Modelling steady-state & light-induced responses of g_s_

Initial steady-state stomatal conductance at 100 μmol m^-2^ s^-1^ (g_s,100_) ranged from 0.016 to 0.032 mol m^-2^s^-1^ (Fig. 2a, Supplemental Table S1). Following the step increase in light to 1000 μmol m^-2^ s^-1^ for 90 min average g_s,1000_ ranged between 0.13 and 0.15 mol m^-2^s^-1^ (Fig. 2a, Supplemental Table S1). The differences observed at both g_s,100_ and g_s,1000_ were not significant between genotypes (Fig. 2a, Supplemental Table S1).

**Fig. 2.**
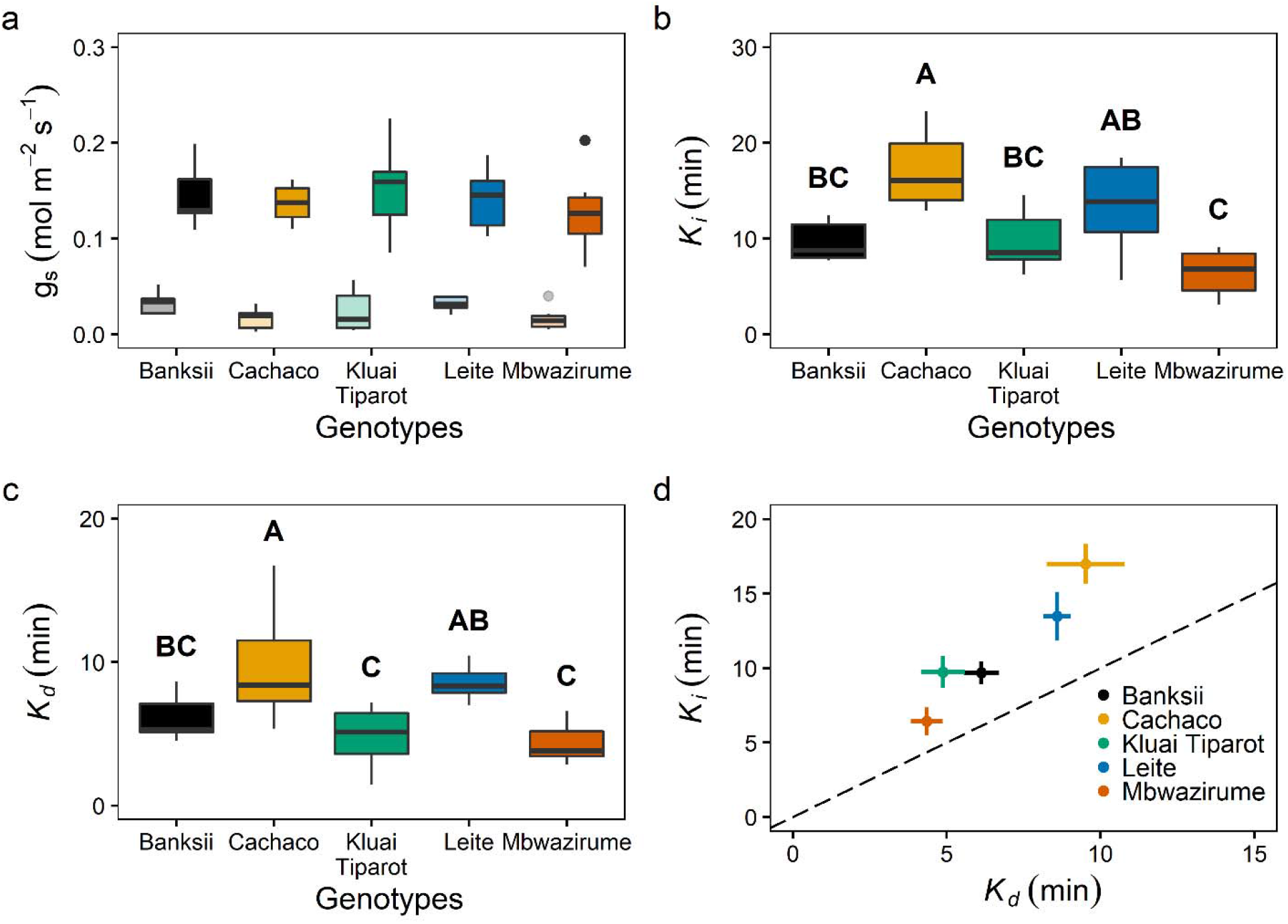
Modelled steady-state and light-induced variables of the stomatal conductance (g_s_) response to a step increase and decrease in light intensity between 100 and 1000 μmol m^-2^ s^-1^ for five different banana genotypes (n = 7-8). (a) Steady-state g_s_ at 100 (g_s,100_ faded colours) and 1000 μmol m^-2^ s^-1^ (g_s,1000_ bright colors). (b) Time constant of g_s_ increase (*K_i_*) for different genotypes. Different letters indicate significant differences between genotypes (P < 0.05; A>B>C). (c) Time constant of g_s_ conductance decrease (*K_d_*) for different genotypes. Different letters indicate significant differences between genotypes (P < 0.05; A>B>C). (d) Significant correlation between *K_i_* and *K_d_* (R^2^ = 0.41, P < 0.001). *K_i_* was significantly higher than *K_d_*. The dashed line shows the 1:1 line. Points and error bars represent mean ± SE (n = 7-8).

The speed of g_s_ increase varied strongly between the *Musa* genotypes and the modelled variables differed significantly (Fig. 2b-c, Supplemental Table S1). The genotype with the slowest g_s_ increase, Cachaco, had an average time constant *K*_i_ of 17 min, while the fastest genotype, Mbwazirume, had a *K*_i_ of 6.4 min (Fig. 2b, Supplemental Table S1). The speed of the decrease in g_s_ (*K*_d_) was also genotype-dependent (Fig. 2c, Supplemental Table S1). *K*_d_ was about 2 fold higher in Cachaco (9.5 min) than in Mbwazirume (4.4 min). Across all genotypes, *K*_i_ was significantly correlated with *K*_d_ (R^2^ = 0.41, P < 0.001; Fig. 2d & Supplemental Fig. S1). However, the decrease in g_s_ was significantly faster than the increase (P < 0.001). The maximal slope of g_s_ increase and decrease (*Sl*_*max*,i_ and *Sl*_*max*,d_) was significantly correlated with the time constant *K* (R^2^ 0.52 and 0.49 for g_s_ increase and decrease respectively, P < 0.001; Supplemental Fig. S1). During light-induced stomatal opening comparable differences across genotypes were present in *Sl*_*max*,i_ as in *K*i. The lowest *Sl*_*max*,i_ values were observed for the genotype Cachaco and the highest values for Mbwazirume (Supplemental Fig. S2, Supplemental Table S1). *Sl*_*max*,d_ was highest for the genotype Kluai Tiparot, while Leite showed the lowest *Sl*_*max*,d_ (Supplemental Fig. S2, Supplemental Table S1). Analogous to the opening and closing time constant, the absolute slope of closing was significantly higher than the opening slope (P < 0.001).

### Impact of stomatal opening and closing speed on *A*

The speed of the increase in g_s_ following a step-change in PPFD from 100 to 1000 μmol m^-2^ s^-1^ strongly determines CO_2_ uptake during this period. The time to reach 95% of maximum *A* was greater than 30 min for almost all genotypes and differed significantly between Cachaco (51.8 min) and the genotypes Mbwazirume (30.3 min) and Banksii (29.5 min) (Fig. 3a, Supplemental Fig. S3). The durations of *A* limitation by g_s_ was significantly correlated with the time constant for g_s_ increase (*K*_i_), confirming the impact of stomatal limitation on *A* (P < 0.001, R^2^ =0.67, Supplemental Fig. S1). The percentage limitation of *A* by g_s_ was significantly higher in Cachaco (20.7 %) compared to the genotypes Mbwazirume (10.2 %), Leite (10.3 %) and Banksii (8.5 %) (Fig. 3b).

**Fig. 3.**
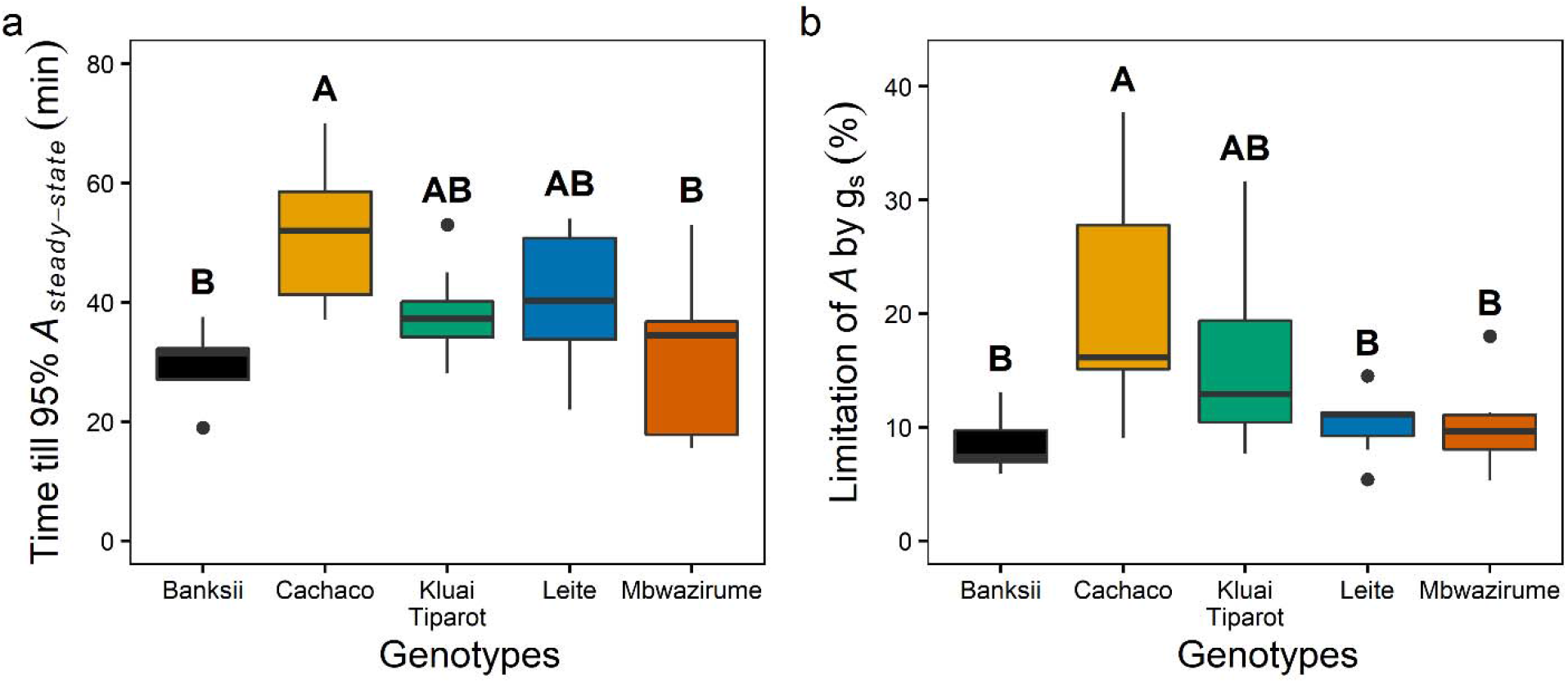
Limitation of steady-state photosynthesis (*A*) by slow stomatal conductance (g_s_) increase after the increase in light intensity from 100 to 1000 μmol m^-2^ s^-1^. (a) Time to reach 95 % of the steady-state *A* for five different banana genotypes. (b) Percentage limitation of *A* by slow g_s_ increase after the increase in light intensity. Different letters indicate significant differences between genotypes (P < 0.05; n = 7-8; A>B).

### _i_WUE response to step-changes in light intensity

The step increase in light intensity induced an initial increase in *A* that was relatively larger than the increase in g_s_. These responsiveness differences increased _i_WUE, reaching the maximum _i_WUE (_i_WUE_max_) during the light period in all cases within 6.5 min (Supplemental Fig. S4). _i_WUE_max_ was reached earlier than steady-state *A*, showing that the maximal _i_WUE and maximal *A* do not coincide. After reaching its maximal value _i_WUE decreased as both g_s_ and *A* gradually increased (Supplemental Fig. S4). _i_WUE only stabilized when both *A* and g_s_ reached steady-state. The genotype Cachaco had a significantly higher cumulative _i_WUE during the high light period compared to Mbwazirume (Supplemental Fig. S5). The cumulative _i_WUE was significantly correlated with the time constant *K*_i_ and *Sl*_*max*,i_ with slower g_s_ responses resulting in higher _i_WUE (R^2^ 0.19 & 0.41, P < 0.01, Supplemental Fig. S1). The reduction in light intensity from 1000 to 100 μmol m^-2^ s^-1^ instantaneously lowered _i_WUE as *A* immediately declined because of light limitation (Supplemental Fig. S4). The cumulative _i_WUE during this low light period was significantly higher in Kluai Tiparot, than in Leite (Supplemental Fig. S5). The cumulative _i_WUE was significantly correlated to the stomatal closing variables *K*_d_ and *Sl*_*max*,d_ with faster g_s_ responses resulting in higher _i_WUE (R^2^ 0.35 & 0.31, P < 0.001, Supplemental Fig. S1).

### Stomatal anatomy

Banana has elliptical-shaped guard cells surrounded by four to six subsidiary cells (Rudall et al., 2017). Abaxial stomatal density, stomatal length, guard cell size and subsidiary cell size were quantified from the leaf part enclosed in the gas exchange cuvette and significant differences between genotypes were observed (Supplemental Fig. S6). Overall these anatomical characteristics were not correlated with any of the modelled light-induced g_s_ kinetics (Fig. 4, Supplemental Fig. S1). However, several correlations between anatomy and g_s_ kinetics were significant if the genotype Cachaco with lowest g_s_ rapidity was not considered (Fig. 4). In this case, stomatal density was significantly correlated with the time constant *K* as well as the maximum slope of g_s_ response *Sl_max_* during both stomatal opening and closing (P < 0.01; R^2^ ranging between 25 and 46 %).

**Fig. 4.**
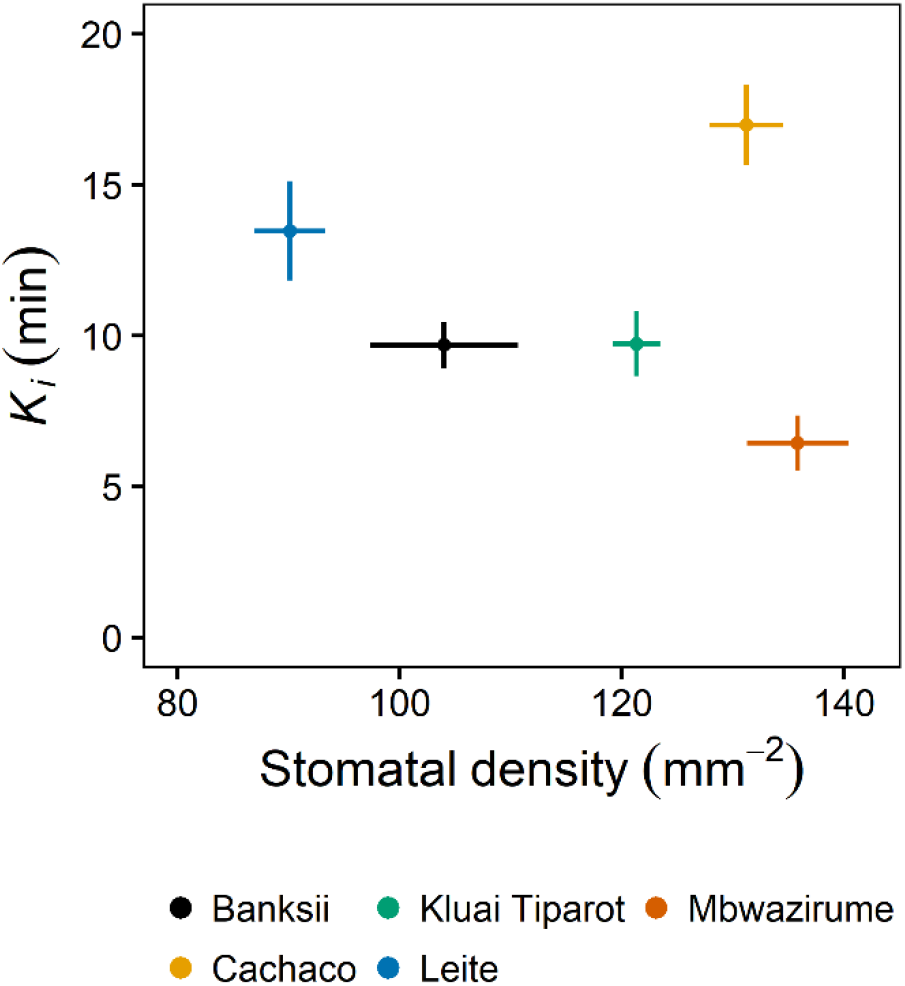
Relation between abaxial stomatal density and the time constant describing the speed of stomatal conductance increase (*K_i_*) after the light intensity increase from 100 to 1000 μmol m^-2^ s^-1^. There was no significant correlation, caused by the outlying genotype Cachaco. Points and error bars represent mean ± SE (n = 7-8).

### Whole-plant transpiration response at dawn

The significant differences at leaf level observed between the two extreme genotypes Cachaco and Mbwazirume were confirmed at the whole-plant level. After the onset of light in the morning, transpiration rate increased significantly earlier in Mbwazirume compared to Cachaco (Fig. 5, P < 0.001). In Mbwazirume a significant increase in transpiration rate was only observed after *c*. 22 min, while in Cachaco this was only after 35 min (Fig. 5a-b). These results were also reflected in the transpiration rate before and after dawn. The whole plant transpiration rate did not differ significantly between both genotypes pre-dawn, but after the onset of light, the transpiration rate was significantly higher in Mbwazirume (Fig 5c).

**Fig. 5.**
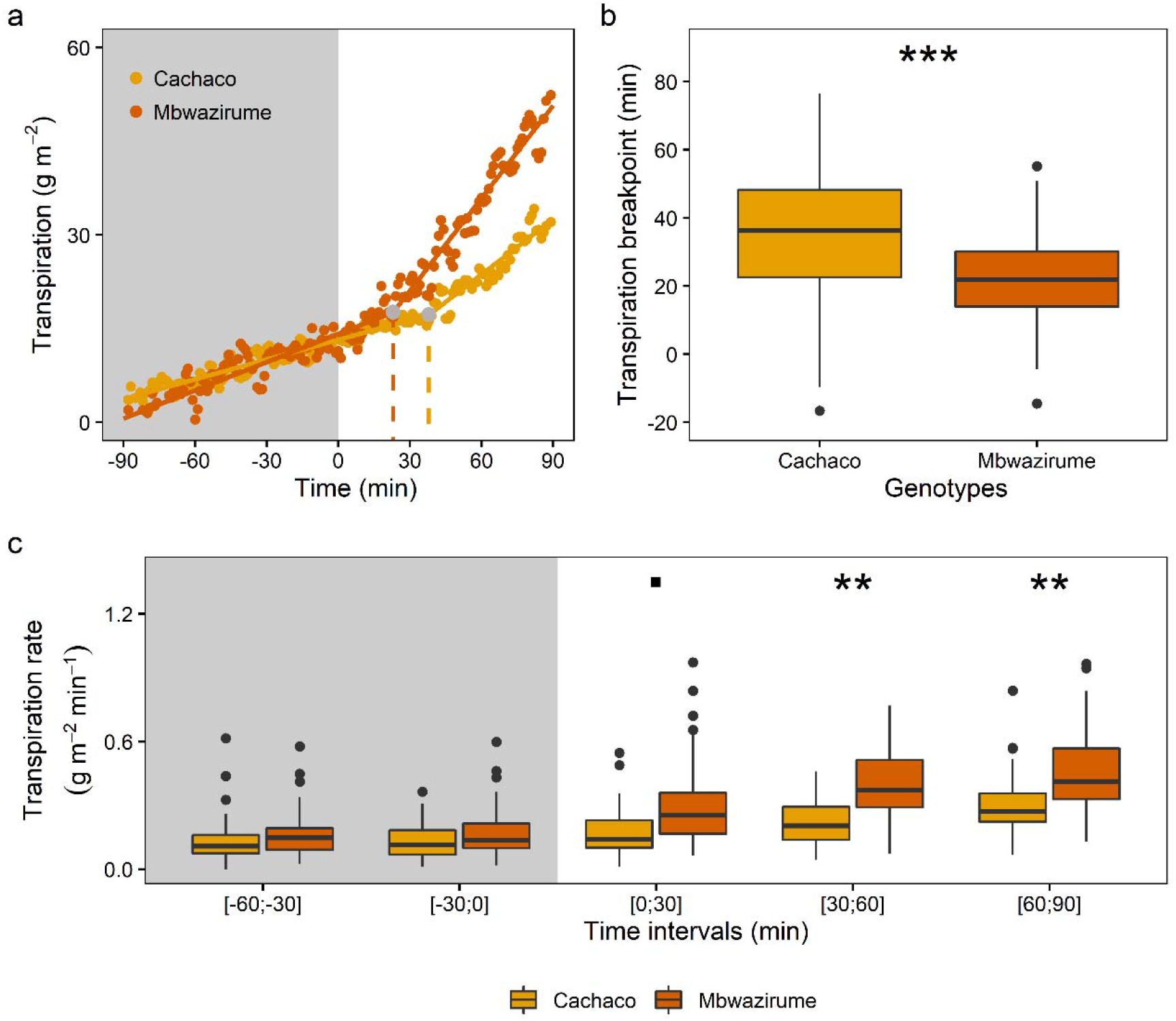
Gravimetric transpiration rate analysis of genotypes Cachaco and Mbwazirume at dawn. (a) A breakpoint was identified in whole-plant transpiration after the onset of light in the morning. The transpiration profiles and breakpoints of a representative plant of each genotype on a selected day are shown. (b) The timing of the breakpoint in transpiration after dawn differed significantly between the genotype Cachaco and Mbwazirume (P < 0.001). The breakpoint indicating an increase in transpiration rate was on average after 22 min in Mbwazirume and 35 min in Cachaco. Only data with significant segmented regression (P < 0.05) and positive slopes were maintained (87 data points for Cachaco and 96 for Mbwazirume). (c) Transpiration rate after dawn increased faster in Mbwazirume compared to Cachaco. Before dawn transpiration rates did not differ significantly, while within 30 min after dawn, transpiration rate differed significantly (76 datapoints per per time range for Cachaco and 79 for Mbwazirume). Grey areas indicate the time before dawn. ‘.’ for P < 0.1, * for P <0.05, ** for P <0.01, *** for P <0.001.

### Impact of diurnal light fluctuations on g_s_, *A* and _i_WUE

To evaluate the impact of g_s_ kinetics on diurnal *A* and _i_WUE, plants were subjected to fluctuating light intensities and phenotyped over an entire diurnal period. Similar to the transpiration rate measured at the whole-plant level, the morning increase in g_s_ at leaf-level under gradually increasing light intensity was faster in Mbwazirume compared to Cachaco (Fig. 6a). The time constant for the g_s_ increase (*K*_i_) was significantly higher in Cachaco (P < 0.005, Fig. 6b). However, the faster increase of g_s_ in Mbwazirume, did not result in increased *A* (Fig. 6c). Maximum potential *A* values at specific light intensities were determined from light response curves and compared to those measured under the diurnal conditions. Under the light intensities experienced in the morning, maximum *A* values were achieved, indicating there was no g_s_ limitation under these light-limiting conditions (Fig. 7). Because of the similar *A* with lower g_s_ during the morning, the mean _i_WUE was significantly higher in Cachaco (P < 0.05, Fig. 6d).

**Fig. 6.**
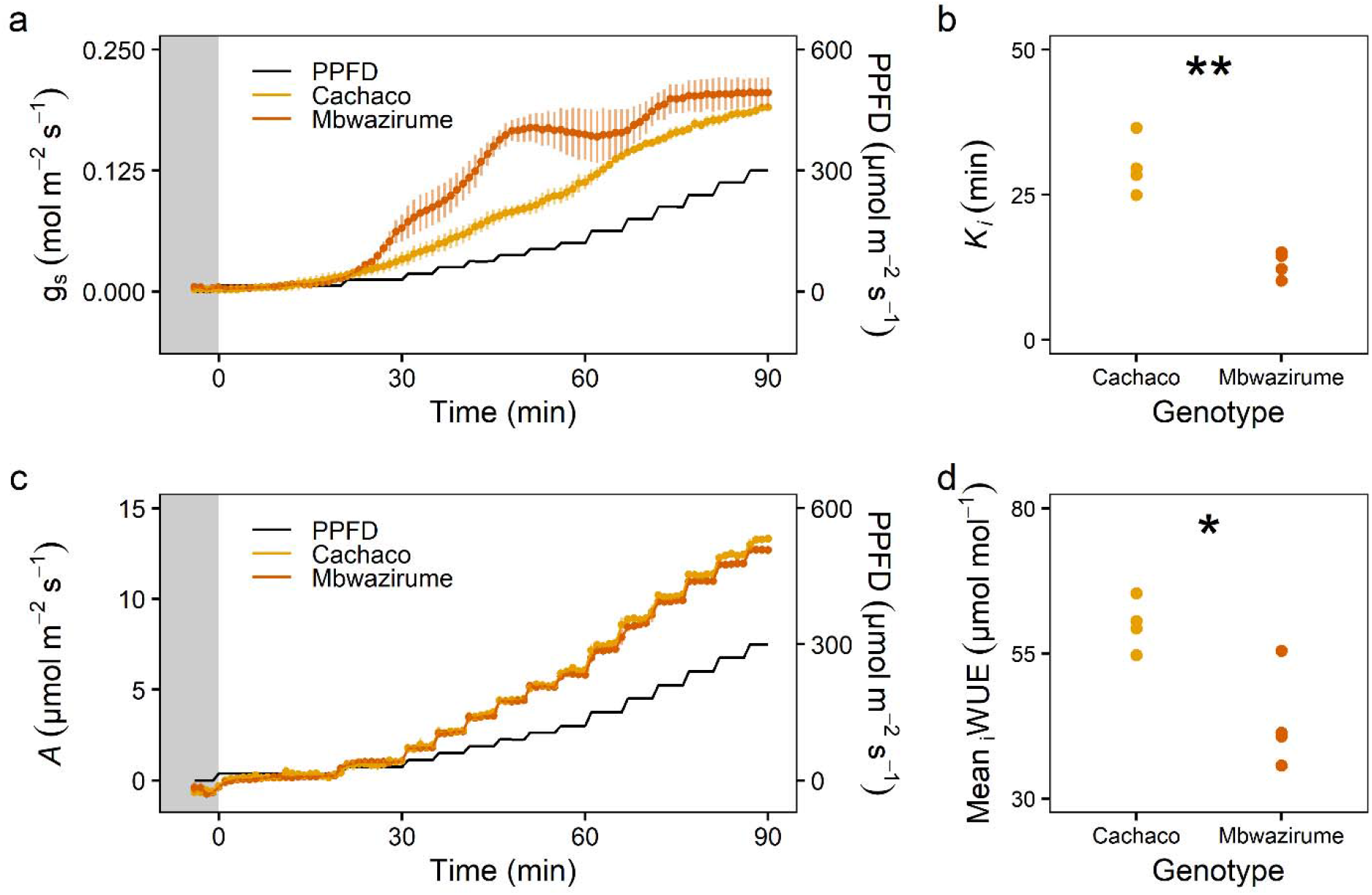
Morning response of stomatal conductance (g_s_) and photosynthesis (*A*) of the genotypes Cachaco and Mbwazirume. (a) Time course of the g_s_ response to a gradual increase in light intensity at dawn (black line). Data are the mean ± SE (n = 4). (b) The time constant of g_s_ increase (*K_i_*) during the first 90 min after dawn was significantly higher in Cachaco. (c) The difference in g_s_ rapidity at dawn did not result in different *A* between both genotypes. Data represents the mean ± SE (n = 4). (d) The mean intrinsic water use efficiency (_i_WUE) during the first 90 min after dawn was significantly higher in Cachaco compared to Mbwazirume. The grey area indicates the time before dawn. PPFD, Photosynthetic Photon Flux Density. *for P <0.05, ** for P <0.01.

Throughout the day, g_s_ kinetics were in most cases significantly faster for the genotype Mbwazirume compared to Cachaco (Fig. 8a), again confirming the previously observed kinetics (Fig. 2, Fig. 5). However, under fluctuating light conditions g_s_ kinetics were dependent on the magnitude of light intensity change, g_s_ values prior to the light intensity change and the time of the day (Fig. 8a). During the afternoon there is a setback in kinetics: the absolute g_s_ and the g_s_ responses to light are damped, strongly limiting *A* (Fig. 7, Fig. 8). The limitation of *A* by g_s_ in the afternoon was three times higher in Cachaco (52.6 %) compared to Mbwazirume (17.5 %) (Fig. 7, Fig. 9d). The reduction of g_s_ in the afternoon resulted in a significantly lower average diurnal g_s_ (Fig. 9a) which translated into a greater diurnal _i_WUE in Cachaco compared to Mbwazirume (Fig. 8c, Fig. 9c).

**Fig. 7.**
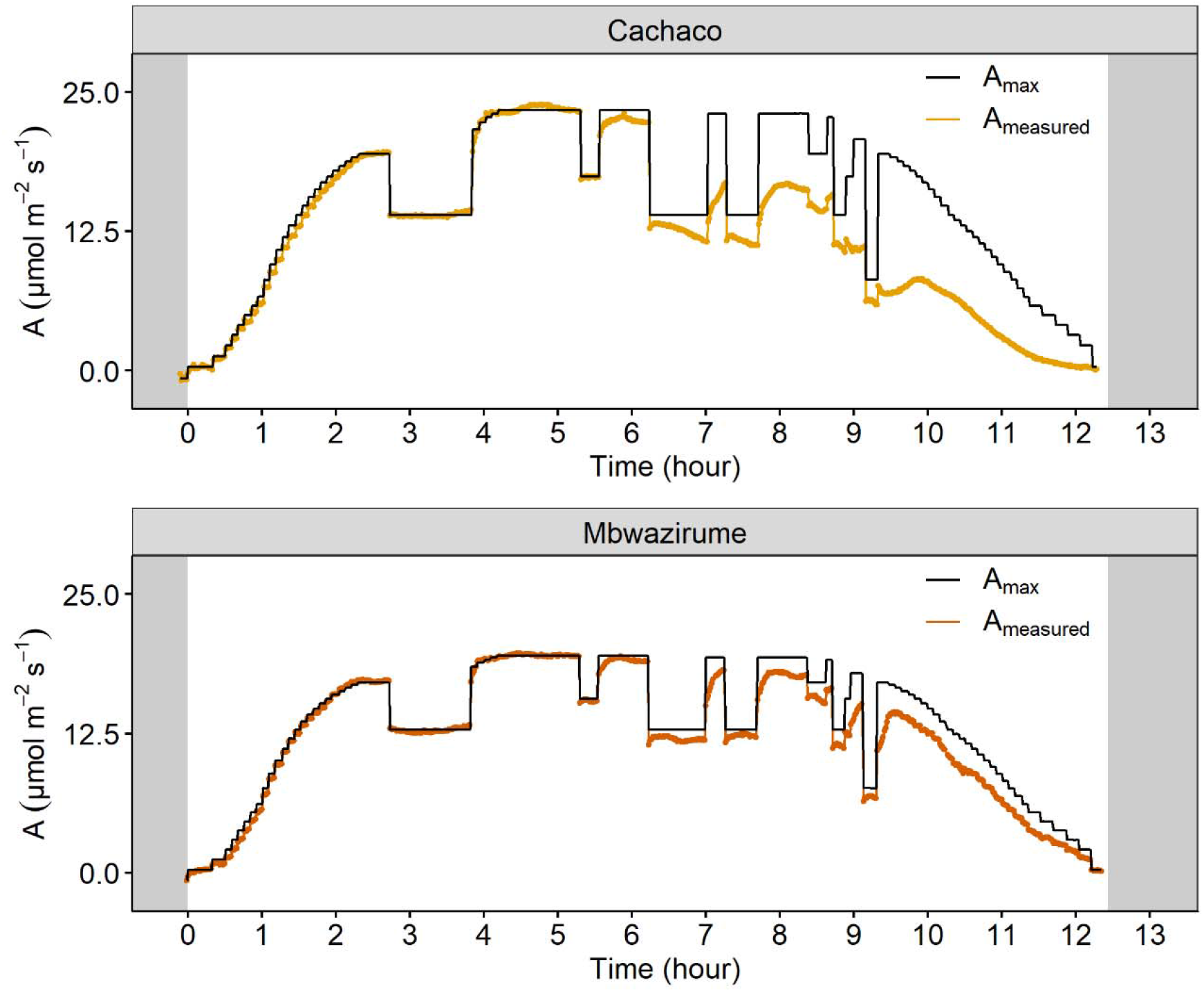
Diurnal time course of measured photosynthesis (*A_measured_*) and maximal photosynthesis (*A_max_*, black line) under fluctuating light conditions for a representative individual of Mbwazirume and Cachaco. The *A_max_* at each light intensity was determined by a modelled light response curve. The nonrectangular hyperbola-based model of Prioul & Chartier (1997) was optimized as described by Lobo et al. (2013). Grey areas indicate times of darkness.

**Fig. 8.**
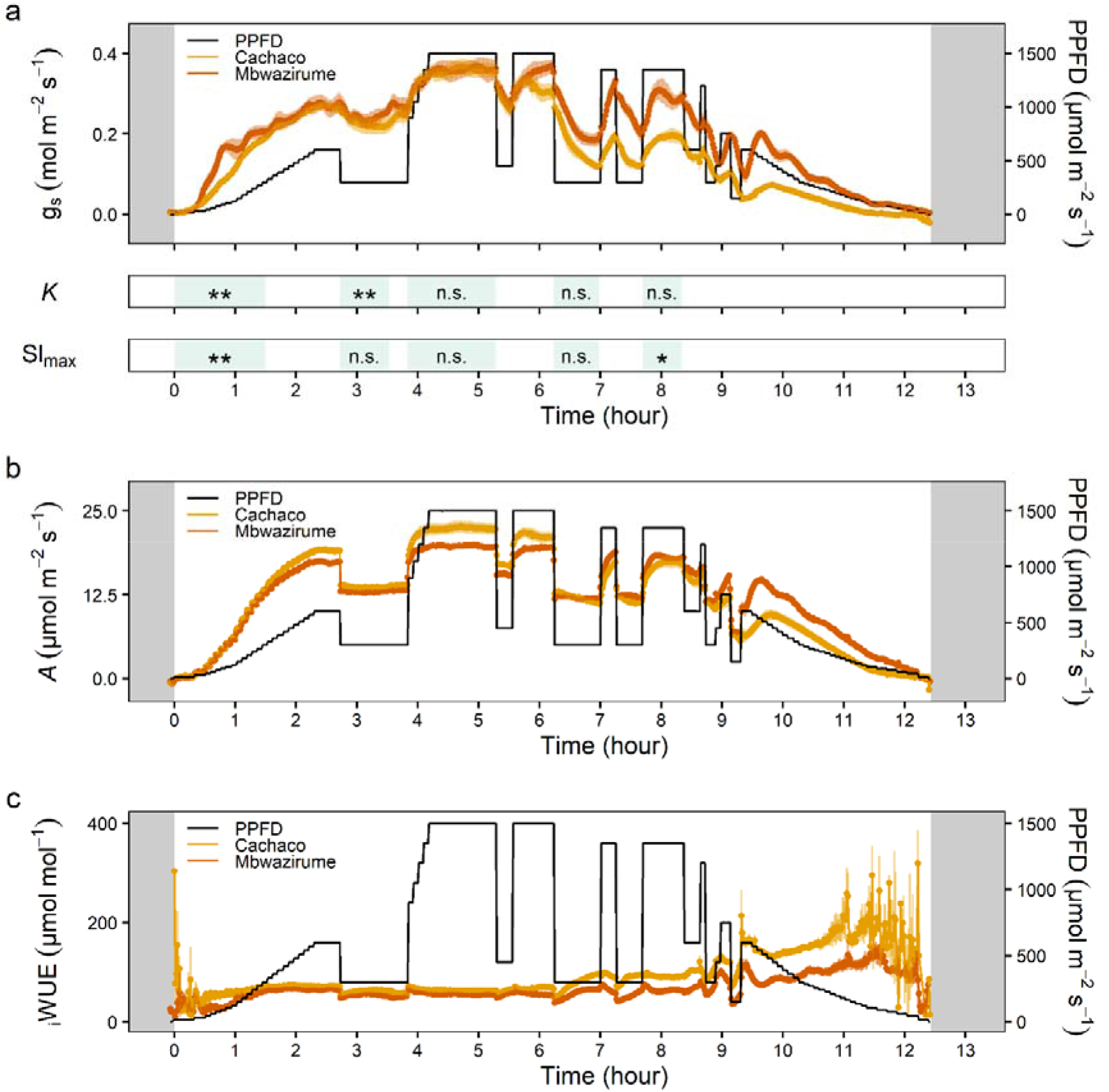
Diurnal time course of (a) stomatal conductance (g_s_), (b) photosynthesis (*A*) and (c) intrinsic water use efficiency (_i_WUE) of the genotypes Mbwazirume and Cachaco. The light intensity fluctuated throughout the day, ranging up to 1500 μmol m^-2^ s^-1^ (black line). The significance of the time constant of g_s_ increase or decrease (*K*) and the maximal slope of g_s_ increase or decrease (*Sl_max_*) is shown. Throughout the day, g_s_ kinetics were faster for the genotype Mbwazirume compared to Cachaco, but differences were dependent on the target light intensity, the magnitude of change, the g_s_ prior to the intensity change and the time of the day. *for P <0.05 and ** for P <0.01 for faster g_s_ rapidity in Mbwazirume compared to Cachaco. Grey areas indicate times of darkness. Green areas indicate the analyzed time frame of the g_s_ rapidity response. PPFD, Photosynthetic Photon Flux Density. Data are the mean ± SE (n = 4).

**Fig. 9.**
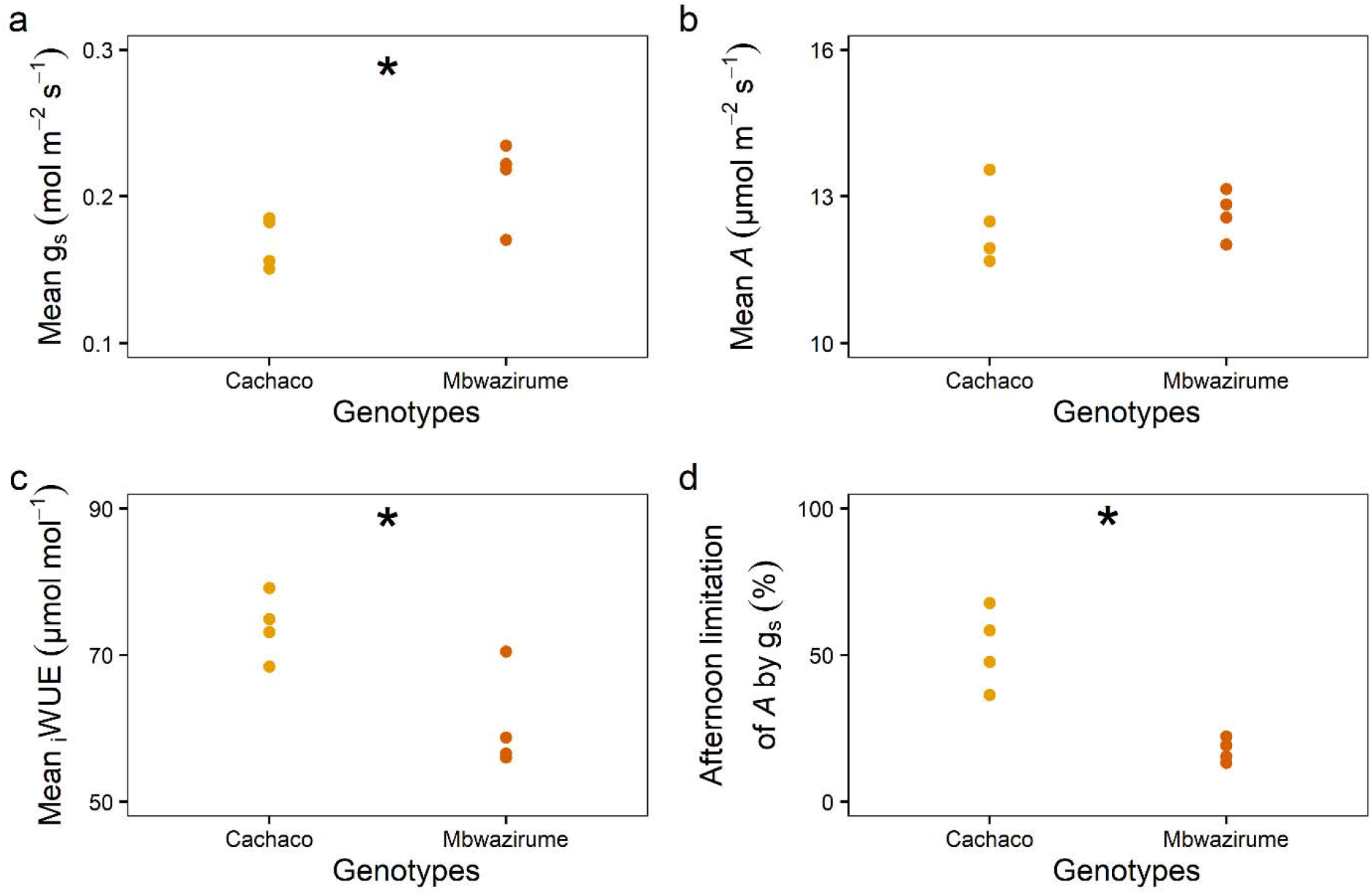
Average diurnal (a) stomatal conductance (g_s_), (b) photosynthesis (*A*) and (c) intrinsic water use efficiency (_i_WUE) of the genotypes Mbwazirume and Cachaco under fluctuating light intensity illustrated in Fig. 7. (d) The percentage limitation of *A* by g_s_ during the afternoon (> 6 hours after light onset).* for P < 0.05.

## Discussion

### Stomatal behaviour greatly limits photosynthesis in banana

Step-changes in light intensity have shown to induce an uncoupling of *A* and g_s_ in many of species (Barradas and Jones, 1996; Lawson and Blatt, 2014; McAusland et al., 2016; Faralli et al., 2019a). However, all *Musa* genotypes maintain a tight coupling between *A* and g_s_ following a step increase in light intensity (Fig. 1). This indicates a strong stomatal control of photosynthesis and overall high limitation of *A* by g_s_ with an average limitation of about 13 % (Fig. 3). This behaviour shows that banana strongly controls stomatal aperture, resulting in water conservation at the expense of potential carbon gain. This prioritizing of water conservation in banana can be explained by its intrinsic need to maintain a high leaf water potential (Turner and Thomas, 1998).

### Diversity in light-induced stomatal responses

Stomatal responses to changes in light intensity have been shown to vary at an inter- and intra-specific level (Vico et al., 2011; Drake et al., 2012; McAusland et al., 2016; Qu et al., 2016; Durand et al., 2020; De Souza et al., 2020). A higher steady-state g_s_ has been linked with faster light-induced g_s_ responses (Drake et al., 2012; Kaiser et al., 2016; McAusland et al., 2016; Wachendorf and Küppers, 2017). Although the differences observed in steady-state g_s_ values between banana genotypes were not significant, their g_s_ kinetics differed strongly (Fig. 2). These results suggest that other factors such as stomatal anatomy, hydraulic conductance and membrane transporters are involved in determining the rapidity of changes in g_s_.

Within the *Musa* family we observed significant differences in the speed of increase and decrease in g_s_ (Fig. 2b,c). Differences across genotypes were not explained by genomic constitution, which is in agreement with the wide diversity of transpiration phenotypes observed irrespective of their genomic constitution (van Wesemael et al., 2019). Consistent with previous works in other species (Vico et al., 2011; McAusland et al., 2016; Faralli et al., 2019a), the speed of g_s_ increase and decrease were significantly correlated (Fig. 3d). This correlation could suggest that the genotypic differences in net solute flux across guard and tonoplast membranes are the main bottlenecks and are maintained in both opening and closing response. Differences in these fluxes could be attributed to differences in the number or activity of such membrane transporters. Decreases in g_s_ were faster than opening in all *Musa* genotypes (Fig. 3d), which is not the case for all crops (McAusland et al., 2016; Qu et al., 2016). The faster g_s_ closure again indicates that *Musa* prioritises water conservation over maximization of carbon uptake.

The two most extreme genotypes Cachaco and Mbwazirume, with the slowest and fastest g_s_ responses respectively, also showed at the whole-plant level, differences in light-induced transpiration rate (Fig. 5). This finding suggests that despite possible differences in g_s_ control of water loss at different locations in the leaf (Matthews et al., 2017) and across leaves of different age (Urban et al., 2008) genotype-specific responses are still maintained. Leaf level measurements of g_s_ kinetics are thus in line with whole-plant responses. To our knowledge, this is the first report confirming stomatal light sensitivity and kinetics at the whole-plant level. The genotype-specific difference in light-induced transpiration responses at dawn was validated at the leaf level with g_s_ increasing faster in Mbwazirume under gradually increasing light intensity (Fig. 6a). This faster g_s_ increase in Mbwazirume did not result in higher *A*, indicating that at dawn, under low light intensities, g_s_ was not limiting *A* and was higher than necessary for maximal *A* (Fig. 6 & Fig. 7). These results demonstrate that the impact of g_s_ kinetics on *A* and _i_WUE depend on the time of the day and the light conditions. The uncoupling of g_s_ and *A* under realistically increasing light conditions at dawn was not beneficial for carbon uptake. Gosa et al. (2019) called this period after dawn in tomato the golden hour because in dry climates it is the time of the day with the highest g_s_. Later in the day, VPDs become too high, restricting g_s_ (Gosa et al., 2019). Breeding for an even higher g_s_ during this golden hour was suggested to improve plant productivity. However, care must be taken to breed for an improved morning CO_2_ uptake, rather than for a high g_s_ with associated uncoupling of *A* and g_s_. Although the absolute water loss resulting from excessive morning g_s_ might be relatively low because of low evaporative demands at dawn (Chaves et al., 2016), it may lead to a crucial decrease in overall plant water status.

Despite the confirmed genotypic differences in stomatal kinetics, the impact of g_s_ kinetics on *A* and _i_WUE before noon hardly differed between the genotypes Cachaco and Mbwazirume under field-mimicking light conditions (Fig. 7, Fig. 8). This could be explained by lower amplitudes of light switches compared to a single step-change in light intensity and/or g_s_ values not being at steady-state prior to changing light intensity. The genotype-specific speed of the g_s_ response observed under a single step-change in light intensity did not explain the diurnal _i_WUE, indicating that g_s_ kinetics only partially affect diurnal water use efficiency and carbon gain (Fig. 9b,c). The absolute g_s_ and the g_s_ responses to light decreased strongly in the afternoon, and this effect was more pronounced in the genotype Cachaco (Fig. 7, Fig. 8a). The three times higher afternoon g_s_ limitation of *A* in the genotype Cachaco compared to Mbwazirume, resulted in a significant higher diurnal _i_WUE (Fig. 9c,d). The genotype Cachaco with the slowest g_s_ kinetics thus achieved the highest _i_WUE, showing that not only g_s_ speed but also the diurnal pattern determines the overall water use efficiency and carbon gain. We show for that under fluctuating light conditions this intrinsic diurnal pattern of absolute g_s_ decrease and g_s_ light responsivity reduction is decisive for diurnal _i_WUE (Fig. 9c).

### Impact of stomatal anatomy on responses

Stomatal density as well as the size have been reported to affect g_s_ kinetics (Hetherington and Woodward, 2003; Drake et al., 2012). However, McAusland et al. (2016) and Faralli et al. (2019a) reported no or only a weak inter- and intra-specific correlation between stomatal anatomy and light-induced g_s_ kinetics. We confirmed that stomatal density and size were not correlated with the g_s_ kinetics (Fig. 4, Supplemental Fig. S1). Remarkably, the genotype with the slowest increase in g_s_, Cachaco had the second highest density and the smallest stomata. Without this genotype a significant correlation between density and the speed of g_s_ increase and decrease was observed (Fig. 4). This exception proves that the surface-to-volumes ratios are not always directly related to stomatal speed as this assumes uniform ion transport activity per surface area (Lawson and Blatt, 2014).

### Conclusion

Our findings prove that there is diversity in g_s_ rapidity to light within closely related genotypes. The priority of banana for water saving is shown by strong stomatal control of *A* and faster decrease in g_s_ than increase. The observed diversity in g_s_ rapidity was not related to stomatal density or subsidiary cell sizes and therefore suggests that variation is driven by functional components rather than anatomy. We show here for the first time that the g_s_ rapidity observed at the leaf level can also be found at the whole-plant level. However, under fluctuating light conditions, g_s_ rapidity is only one of the many physiological factors determining overall plant water use efficiency and carbon gain.

## Materials & methods

### Experiment 1: Leaf gas exchange response to a step-change in light intensity

#### Plant material and growth conditions

Banana plants were obtained through the International Musa Transit Center (ITC, Bioversity International), hosted at KU Leuven, Belgium. Plants of five genotypes from different subgroups were selected: Banksii (subgroup Banksii, AA genome, ITC0623), Cachaco (Bluggoe, ABB genome, ITC0643), Kluai Tiparot (Kluai Tiparot, ABB genome, ITC0652), Leite (Rio, AAA genome, ITC0277) and Mbwazirume (Mutika-Lujugira, AAA genome, ITC1356). Plants were grown in 800 ml containers filled with peat-based compost (Levingtons F2S, UK) under 350 μmol m^-2^ s^-1^ Photosynthetic Photon Flux Density (PPFD) in a 12 h: 12 h light: dark cycle with temperature and relative humidity at 26 ± 1 °C and 70 ± 10 %, respectively. Plants were well-watered and starting from week three a Hoagland nutrient solution was added. Measurements were performed when plants were fully acclimated and seven weeks old.

#### Leaf gas exchange measurements

Photosynthetic rate (*A*) and stomatal conductance to water (g_s_) were measured every 30 s on the middle of the second youngest fully developed leaf using a LI-6400XT infrared gas analysis and dew-point generator model LI-610 (LI-COR, USA). Light was applied by an integrated LED light source. The leaf cuvette maintained a CO_2_ concentration of 400 μmol mol^-1^, a leaf temperature of 25°C and a VPD of 1 kPa. All measurements were performed before 14:00 h to avoid circadian influences on *A* by g_s_.

#### Stomatal response to a step-change in light intensity

The light intensity was kept at 100 μmol m^-2^ s^-1^ until *A* and g_s_ were stable for 10 min. Once steady-state was reached, light intensity was increased to 1000 μmol m^-2^ s^-1^ for 90 min. Then, light intensity was lowered back to 100 μmol m^-2^ s^-1^ for 30 min.

The increase in g_s_ after the increase in light intensity and the decrease in g_s_ after the decrease in light intensity followed a sigmoidal pattern and was modelled using the non-linear sigmoidal model of Vialet-Chabrand et al. (2013):

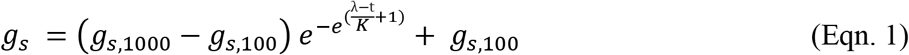

With g_s_ the stomatal conductance at time t, *K* the time constant for rapidity of g_s_ response (min), λ the initial time lag before g_s_ increase or decrease (min), g_s,100_ and g_s,1000_ (mol m^-2^ s^-1^) the steady-state g_s_ at 100 and 1000 μmol m^-2^ s^-1^respectively. Parameter values were estimated for each individual plant using non-linear model optimization in R (V 3.4.3). *K*_i_ indicates the g_s_ increase time constant, *K*_d_ the g_s_ decrease time constant. The maximum slope of g_s_ during opening and closing was calculated and defined as *Sl_max_*. Intrinsic water use efficiency (_i_WUE) was calculated as _i_WUE = *A*/g_s_. Outlying values (0.5 % quantile; _i_WUE < 0 or > 400 μmol mol^-1^) caused by low g_s_ were discarded for plotting. Photosynthesis was considered to be limited by stomatal conductance during the light-induced stomatal opening until 95 % of steady-state *A* was reached (McAusland et al., 2016). The percentage of limitation of *A* by g_s_ was calculated by comparing the measured *A* with the maximal potential *A* without g_s_ limitation according to McAusland et al., (2016):

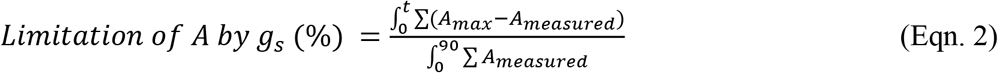

With *A_max_* the value reached at 95 % of steady-state *A, A_measured_* the measured *A* and t the time where 95 % of steady-state *A* is reached.

#### Stomatal anatomy measurements

Stomatal impressions of the abaxial surface of the leaf were made when stomata were completely closed using impression material. Impression were made by applying dental polymer according to the protocol of Weyers & Johansen (1985), followed by covering the polymer with nail varnish and placement on a microscope slide. Stomatal anatomy was quantified using an EVOS digital inverted microscope. Stomatal density was determined in three microscopic field of views of 1.12 mm^2^ captured with a 10 x objective lens (54 to 117 stomata per field of view). Guard cell length (μm), guard cell size (mm^2^) and lateral subsidiary cell size (mm^2^) were determined in three microscopic field of views of 0.07 mm^2^ captured with a 40 x magnification respectively (four to seven stomata per field of view). Measurements were performed in ImageJ software (http://rsb.info.nih.gov/ij).

### Experiment 2: Whole-plant transpiration response at dawn

#### Plant material and growth conditions

For the genotypes Cachaco and Mbwazirume, 12 plants were grown for seven weeks in a greenhouse prior to the experiment. Plants were grown in 10 l containers filled with peat-based compost. At the start of the experiments, the six most homogenous plants per genotype were selected based on leaf area. Weight of each plant was followed by a multi-lysimeter setup of high precision balances (1 g accuracy, Phenospex, Heerlen, NL). The soil was covered by plastic to avoid evaporation and ensure only waterloss trough transpiration. The transpiration rate was calculated by differentiating the raw weight data over time. The soil water content was determined by subtracting the plastic pot weight, the dry soil weight and the plant weight from the total weight measurement. Dry soil weight was calculated as a function of the soil volume (bulk density = 0.2267 g cm^-3^). Leaf area was calculated by weekly top view imaging and modelled over time by a power law function (Paine et al., 2012):

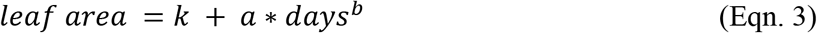

The daily plant weight was estimated from the projected leaf area using genotype-specific correlations (n > 50; R^2^ > 0.94). Plants were watered with a nutrient solution during the night and kept at well-watered conditions. Radiation was collected every 5 min via a sensor (Skye instruments, UK) inside the greenhouse. Supplemental lighting of 14 W m^-^2 at plant level was provided when solar radiation was below 250 W m^-^2 during the daytime. Temperature and relative humidity data were collected using six data loggers (Trotec, DE) registering data every 5 min. The onset of light was defined as the moment when intensity increased above 2 W m^-^2.

### Experiment 3: Impact of diurnal light fluctuations on g_s_, *A* and _i_WUE

#### Plant material and growth conditions

Four plants of the genotypes Cachaco and Mbwazirume were grown in a greenhouse. Plants were grown in 4 l containers filled with peat-based compost and maintained under well-watered conditions. After 8 weeks plants were moved to a growth chamber with relative humidity 70 ± 15 % and temperature 28 ± 2 °C.

#### Leaf gas exchange measurements

*A* and g_s_ were measured every minute on the middle of the second youngest fully developed leaf using a LI-6800 infrared gas analyser (LI-COR, USA). The leaf cuvette maintained a CO_2_ concentration of 400 μmol mol^-1^, a leaf temperature of 28 °C and a VPD of 1 kPa. The light intensity was programmed to fluctuate throughout the day. Plants were placed under adjustable LED panels (LuminiGrow 600R1, Lumini technology Co. Ltd., China) that mimicked light fluctuations inside the LI-6800 leaf cuvette. The g_s_ response was described using the non-linear sigmoidal model of Vialet-Chabrand et al. (2013) (Eqn. 1).

A light response curve with *A* in function of PPFD was modelled for each individual based on *A* values recorded during the first six hours of the day that were not limited by g_s_. The nonrectangular hyperbola-based model of Prioul & Chartier (1977) was optimized as described by Lobo et al. (2013):

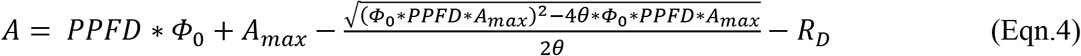

With *A* the photosynthetic rate (μmol m^-2^ s^-1^), PPFD the Photosynthetic Photon Flux Density (μmol m^-2^ s^-1^), Φ_0_ the quantum yield at PPFD of 0 μmol m^-2^ s^-1^ (μmol μmol^-1^), *A_max_* the maximum photosynthetic rate (μmol m^-2^ s^-1^), θ the dimensionless convexity factor and R_d_ the dark respiration (μmol m^-2^ s^-1^).

The percentage of limitation of *A* by g_s_ during the afternoon (> 6 hours after light onset) was calculated by estimating the maximal potential *A* without g_s_ limitation and comparing it with the measured *A* (Eqn. 2).

### Statistical analysis and data processing

All data processing and statistical analysis was carried out in R (V 3.4.3). Genotypic differences were tested by applying one-way analysis of variance (ANOVA) with a post hoc Tukey HSD test. Segmented regression was performed on the whole-plant transpiration between −90 and 90 min relative to the onset of light. Only data with significant segmented regression (p-value Davies Test < 0.05, segmented R package) and positive slopes were maintained. Transpiration rate was calculated as the mean water loss every 30 min. Statistical interpretation of results took into account repeated measurements by incorporating a plant-specific factor and a date-specific factor as a random effect in a linear mixed model. The bold middle line in boxplots represents the median The box is confined by the first and third quartile and the whiskers extend to 1.5 times the interquartile distance. Points falling outside the whiskers are considered outliers and plotted as dots.

## Acknowledgement

The authors would like to thank Edwige André for the plant propagation and growth; Hendrik Siongers, Loïck Derette, Simon Costers and Kaat Hebbelinck for technical assistance during plant growth and phenotyping; Silvere Vialet-Chabrand and Phil Davey for their technical assistance during the step-changes in light intensity experiments.

## Supplemental data

**Supplemental Fig. S1** Correlation matrix of gas exchange and stomatal anatomy variables.

**Supplemental Fig. S2** Maximum slope of stomatal conductance response (*Sl_max_*) to an increase in light intensity from 100 μmol m^-2^ s^-1^ to 1000 μmol m^-2^ s^-1^ and to a decrease in light intensity from 1000 μmol μmol m^-2^ s^-1^ to 100 μmol m^-2^ s^-1^

**Supplemental Fig. S3** Increase in photosynthesis (*A*) after increasing the light intensity from 100 μmol m^-2^ s^-1^ to 1000 μmol m^-2^ s^-1^.

**Supplemental Fig. S4** Normalized response of intrinsic water use efficiency (_i_WUE) to a step increase and decrease in light intensity from 100 to 1000 μmol m^-2^ s^-1^ and back.

**Supplemental Fig. S5** Cumulative intrinsic water use efficiency (_i_WUE) after the increase in light intensity from 100 to 1000 μmol m^-2^ s^-1^ and the decrease to 100 μmol m^-2^ s^-1^ afterwards.

**Supplemental Fig. S6** Stomatal density, stomatal length, guard cell size, subsidiary cell size and proportion of subsidiary cells of the five banana genotypes.

**Supplemental Table S1** Modelled steady-state and light-induced variables of the stomatal conductance (g_s_) response to a step increase and decrease in light intensity from 100 to 1000 μmol m^-2^ s^-1^ for five different banana genotypes (Musa spp.).

